# In vivo analysis of the contribution of proprotein convertases to the processing of FGF23

**DOI:** 10.1101/2021.01.15.426882

**Authors:** Omar Al Rifai, Delia Susan-Resiga, Rachid Essalmani, John W.M. Creemers, Nabil G. Seidah, Mathieu Ferron

**Affiliations:** Unité de recherche en physiologie moléculaire, Institut de Recherches Cliniques de Montréal, Montréal, Québec H2W 1R7, Canada; Programme de biologie moléculaire, Université de Montréal, Québec H3T 3J7, Canada; Unité de recherche en biochimie neuroendocrinienne, Institut de Recherches Cliniques de Montréal, Montréal, Québec H2W 1R7, Canada; Department of Human Genetics, KU Leuven, Herestraat 49, B-3000 Leuven, Belgium; Département de Médecine, Université de Montréal, Québec H3T 3J7, Canada; Division of Experimental Medicine, McGill University, Montréal, Québec H3A 1A3, Canada

**Keywords:** FGF23, endoproteolysis, proprotein convertases, Furin, PC5, PACE4, phosphate

## Abstract

Fibroblast growth factor 23 (FGF23) is a hormone secreted from fully differentiated osteoblasts and osteocytes that inhibits phosphate reabsorption by kidney proximal tubules. The full-length (i.e., intact) protein mediates FGF23 endocrine functions, while endoproteolytic cleavage at a consensus cleavage sequence for the proprotein convertases (PCs) inactivates FGF23. Two PCs, furin and PC5, were shown to cleave FGF23 *in vitro* at RHTR_179_↓, but whether they are fulfilling this function in vivo is currently unknown. To address this question we used here mice lacking either or both furin and PC5 in cell-specific manners and mice lacking the paired basic amino acid-cleaving enzyme 4 (PACE4) in all cells. Our analysis shows that furin inactivation in osteoblasts and osteocytes results in a 25% increase in circulating intact FGF23, without any significant impact on serum phosphate levels, whether mice are maintained on a normal or a low phosphate diet. Under conditions of iron deficiency, FGF23 is normally processed in control mice, but its processing is impaired in mice lacking furin in osteoblasts and osteocytes. In contrast, FGF23 is normally cleaved following erythropoietin or IL-1β injections in mice lacking furin or both furin and PC5, and in PACE4-deficient mice. Altogether, these studies suggest that furin is only partially responsible for FGF23 cleavage under certain conditions *in vivo*. The processing of FGF23 may therefore involve the redundant action of multiple PCs or of other peptidases in osteoblasts, osteocytes and hematopoietic cells.

## Introduction

Fibroblast growth factor 23 (FGF23) is a bone-derived hormone (i.e., osteokine) secreted by fully differentiated osteoblasts and osteocytes. FGF23 plays a fundamental role in phosphate and vitamin D metabolism through a direct action on the kidney ^1,2^. In this tissue, its signaling is mediated through interaction with the FGF receptor 1 (FGFR1) and its co-receptor α-klotho ^3^. FGF23 signaling through FGFR1/α-klotho in the kidney proximal tubule suppresses the expression of *Slc34a1* and *Slc34a3*, the genes encoding for the sodium phosphate co-transporters NaPi2A and NaPi2C, thereby increasing urinary phosphate excretion and decreasing serum phosphate levels. In addition, FGF23 reduces the 1,25-dihydroxyvitamin D_3_(calcitriol) level by decreasing expression of *Cyp27b1* which encodes the 25-hydroxyvitamin D-1 alpha hydroxylase and increasing expression of *Cyp24a1* which encodes the 1,25-dihydroxyvitamin D_3_24-hydroxylase ^4–7^. These endocrine functions are mediated through full-length or intact FGF23 (amino acids [aa] 25-251). Endoproteolytic cleavage of FGF23 at the basic motif R_176_HTR_179_↓SA, a consensus cleavage site for proprotein convertases (PCs), releases N-terminal (aa 25-179) and C-terminal (aa 180-251) fragments and inactivates FGF23.

Circulating intact FGF23 levels are tightly regulated in part through various post-translational modifications of amino acids surrounding the PC cleavage site. For instance, the *O*-glycosylation of threonine 178 (T_178_) by the *N*-acetylgalactosaminyltransferase 3 (GalNAc-T3) inhibits FGF23 processing ^8–10^. FGF23 *O-*glycosylation is in turn reduced when the serine 180 (S_180_) is phosphorylated by Fam20c, a kinase residing in the Golgi ^11^. Impaired FGF23 glycosylation caused either by a defective GalNAc-T3 or mutation in FGF23 glycosylation sites, results in an increase in FGF23 processing and subsequent decrease in circulating intact FGF23 levels. These molecular defects lead to familial tumoral calcinosis (FTC), a human disease characterized by hyperphosphatemia and heterotopic calcification ^8–10^. On the other hand, mutations in the FGF23 cleavage site and in the X-linked phosphate-regulating neutral endopeptidase (PHEX) respectively cause autosomal dominant hypophosphatemic rickets (ADHR) and X-linked hypophosphatemic rickets (XLH) that are characterized by hypophosphatemia and osteomalacia due to the increased level of active intact FGF23 ^12,13^.

Iron deficiency, erythropoietin treatment or interleukin-1 beta (IL-1β) stimulate FGF23 production by osteocytes and bone marrow hematopoietic cells ^14–19^. However, this increase in FGF23 production only results in the accumulation of intact FGF23 levels and hypophosphatemia in ADHR patients or in mice with an ADHR-like mutation in *Fgf23*. In wildtype animals or healthy subjects, the same stimuli result in increased circulating levels of cleaved FGF23, revealing the importance of endoproteolysis by PC for the regulation of this hormone.

The proprotein convertases furin and PC5 can cleave FGF23 *in vitro* ^20^, and *FURIN* inactivation in the human osteosarcomas U2-OS cell line impaired the cleavage of ectopically expressed human FGF23 ^11^. Nevertheless, the identity of the enzyme(s) responsible for FGF23 endoproteolysis *in vivo* remains undetermined. In the present study we assessed whether furin and PC5 cleave FGF23 in vivo and affect phosphate metabolism. We showed that unlike what is the case in vitro, furin inactivation in the osteoblastic lineage results only in a modest increase in intact FGF23 under normal or low phosphate diet, but impairs FGF23 cleavage in conditions of iron deficiency. We also show that furin, PC5 and PACE4, another PC expressed in osteoblasts, which all can cleave FGF23 *in vitro*, failed to do so *in vivo* following treatment with erythropoietin or IL-1β. Together, our data suggest that furin, PC5 and PACE4 may not be the main PCs cleaving FGF23 *in vivo*.

## RESULTS

### In vivo inactivation of Furin in osteoblasts and osteocytes partially impairs FGF23 cleavage, but decreases phosphate excretion

To investigate the role of furin in the regulation of FGF23 processing *in vivo*, we generated mice in which the *Furin* gene was specifically inactivated in osteoblasts and osteocytes (i.e., *Furin*^*osb*^*-/-*) by breeding *Furin*^*fl/fl*^ mice with hOCN-Cre transgenic animals which express the Cre recombinase under the control of human osteocalcin promoter ^21^. We previously reported that the *Furin* gene was efficiently and specifically inactivated in osteoblasts and bones in these mice ^22^. Although the precursor of osteocalcin, another bone-derived hormone involved in the control of glucose homeostasis, is not processed in *Furin*^*osb*^*-/-*mice, the same animals do not develop glucose intolerance before 9 months of age ^22^. Thus, to avoid a potential confounding effect of hyperglycemia on FGF23 and phosphate metabolism, all the following experiments were performed on mice aged between 3 and 6 months. When fed a normal chow diet containing ∼0.7% inorganic phosphate (Pi), 4-month-old *Furin*^*osb*^*-/-*female mice displayed a significant 25% rise in circulating intact FGF23 (Figure 1A). This rise was however more moderate than expected in view of in vitro-based studies ^11^ and did not significantly alter the expression of *Slc34a1, Slc34a3, Cyp27b1* or *Cyp24a1* (Figure 1B). Accordingly, *Furin*^*osb*^*-/- mice* maintained normal serum phosphate levels and bone histology did not reveal presence of osteomalacia (Figure 1C and D).

**Figure 1:**
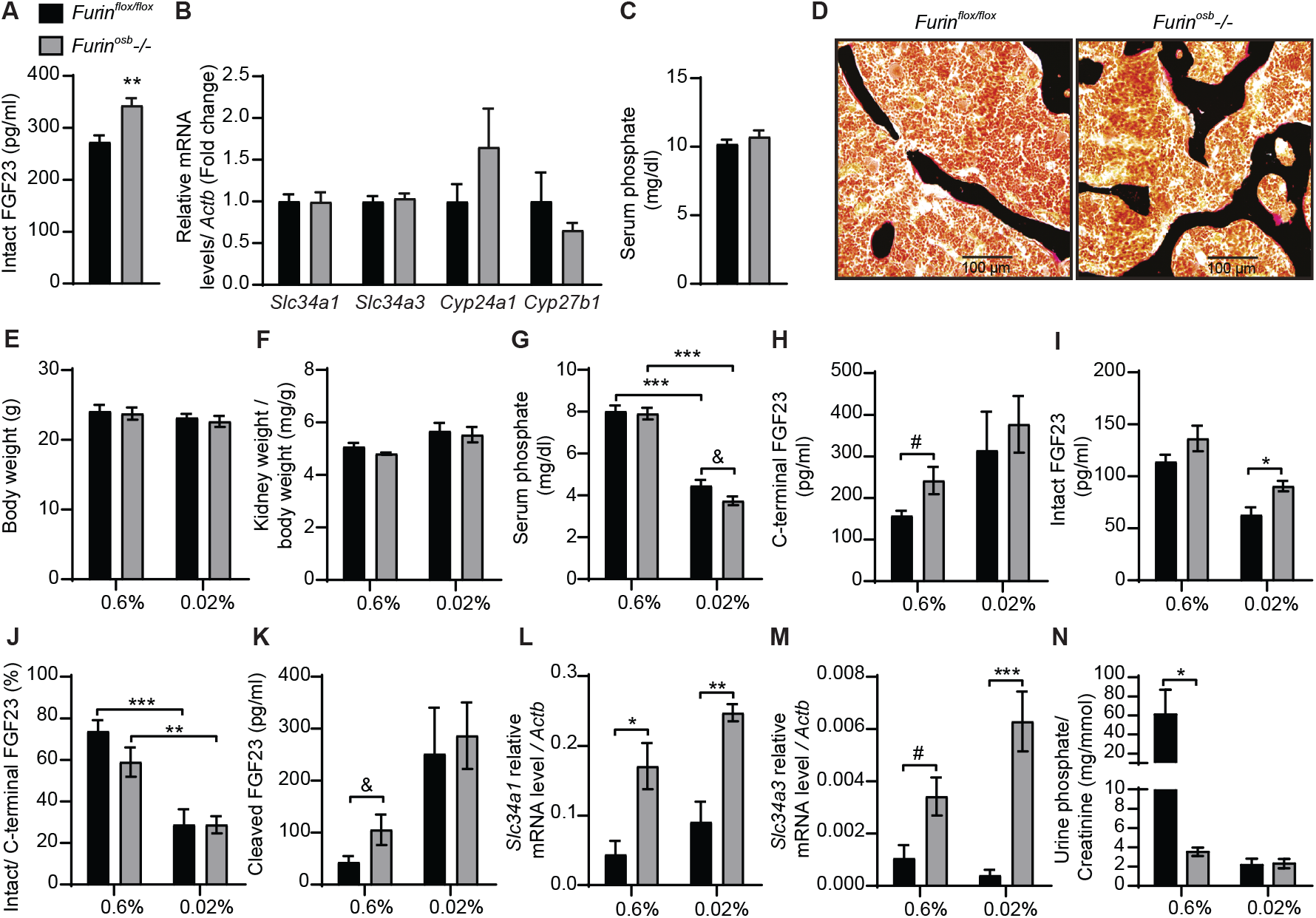
Increased FGF23 level and reduced phosphate excretion in *Furin*^*osb*^*-/-*mice. FGF23 and phosphate levels in in 4-month-old *Furin*^*flox/flox*^ (n=6) and *Furin*^*osb*^*-/-*(n=6) mice fed on normal chow diet **(A-D)**. Intact FGF23 levels in plasma **(A)**. FGF23 target genes expression in kidney **(B)**. Serum phosphate level **(C)**. Von Kossa / van Gieson staining on vertebrae sections **(D)**. FGF23 and phosphate parameters in 6-months-old *Furin*^*flox/flox*^ (n=6) and *Furin*^*osb*^*-/-*(n=6-7) mice fed a normal phosphate diet (0.6%) or a low phosphate diet (0.02%) for one week **(E-N)**. Body weight **(E)**. Kidney weight normalize to body weight **(F)**. Serum phosphate levels **(G)**. Plasma level of C-terminal FGF23 **(H)**, intact FGF23 **(I)**, percentage (%) of intact over C-terminal FGF23 **(J)** and cleaved FGF23 levels **(K)** calculated as (C-terminal FGF23) - (intact FGF23). Expression level in kidney of the sodium phosphate cotransporter *Slc34a1* **(L)** and *Slc34a3* **(M). (N)** Urine phosphate measurements normalized to creatinine. Results represent the mean ± SEM. \**P* < 0.05, \*\**P* < 0.01, and \*\*\**P* < 0.001, by 2-way ANOVA with Bonferroni’s multiple comparisons test or by Student’s t-test (A, C, G, H, K, and M) #: *P*<0.05; # #: *P*<0.01, # # #: *P*<0.001 &: 0.05<*P*<0.1.

Serum phosphate level is a potent regulator of FGF23 and in mice fed a low-phosphate synthetic diet the circulating intact FGF23 levels are reduced, while those of the processed fraction are increased, presumably due to the action of one or several PCs ^23–25^. We therefore next assessed if severe hypophosphatemia would be induced in *Furin*^*osb*^*-/- mice* by reducing their phosphate intake. We found that feeding control and *Furin*^*osb*^*-/- mice* a low-phosphate diet (0.02% Pi) for one week did not affect body weight or kidney weight (Figure 1E and F), but significantly decreased serum phosphate level (Figure 1G). Under the same conditions, *Furin*^*osb*^*-/- mice* tend to have a lower serum phosphate level compared to control mice, although this did not reach statistical significance. *Furin*^*osb*^*-/- mice* on normal diet had >25% increase in intact and C-terminal (total) FGF23, while maintaining normal intact/C-terminal FGF23 ratio compared to control littermates (Figure 1H-J). Under low phosphate diet circulating intact FGF23 was reduced in control mice, but remains significantly higher in *Furin*^*osb*^*-/- mice* (Figure 1I). However, the ratio of intact over C-terminal FGF23 was decreased in both control and *Furin*^*osb*^*-/- mice* under low phosphate diet (Figure 1J). Moreover, the amount of circulating cleaved FGF23 (i.e., C-terminal FGF23 minus intact FGF23) was increased on low phosphate diet regardless of the genotype, suggesting efficient cleavage of FGF23 even in absence of furin (Figure 1K). These results suggest that in conditions of normal or low phosphate intake, processing of FGF23 still occurs in absence of furin in osteoblasts and osteocytes.

Surprisingly, when fed synthetic normal or low phosphate diets, *Furin*^*osb*^*-/- mice* displayed an increased expression in the kidney of the sodium phosphate transporter genes *Slc34a1* and *Slc34a3* compared to control mice, regardless of the level of phosphate in the diet (Figure 1L and M). In addition, *Furin*^*osb*^*-/- mice* fed the normal phosphate diet showed a decrease in urinary phosphate compared to control littermate (Figure 1N). Challenging the mice for one week on the low phosphate diet reduced urinary phosphate in control mice as expected, but not in *Furin*^*osb*^*-/-*mice, as these mice maintained the low urinary phosphate observed on normal phosphate diet (Figure 1N). Together these results show that unlike what was observed in vitro, in both physiological conditions or during decreased phosphate intake, inactivation of *Furin* in the osteoblast lineage has only a modest impact on FGF23 processing. They also suggest that furin may influence phosphate excretion through FGF23-independent mechanisms.

### Furin contributes to FGF23 processing during iron deficiency

It was previously reported that mice carrying an ADHR mutation (R176Q, i.e., loss of P4 Arg in the motif **R**_176_HT**R**_179_↓SA) in the *Fgf23* gene did not develop hypophosphatemia unless they were fed a low-iron diet ^15^. In view of these observations, we induced iron-deficiency in control and *Furin*^*osb*^*-/- mice* by feeding them a low-iron diet in combination with repeated tail bleedings (i.e., ∼70 μl every other week) for 14 weeks, before assessing the impact on FGF23 and phosphate metabolism. Control and *Furin*^*osb*^*-/- mice* fed this diet grew normally (Figure 2A), but both developed iron deficiency as reflected by the decrease in hepcidin (*Hamp*) expression and the increase in transferrin receptor (*Tfrc*) expression in liver (Figure 2B-C). As previously shown ^15^, iron deficient control and *Furin*^*osb*^*-/- mice* displayed an increase in *Fgf23* gene expression in bone (Figure 2D). However, iron-deficiency resulted in increased in circulating C-terminal FGF23 only in control mice, although both genotypes showed an increase in circulating intact FGF23 (Figure 2E-F). These results indicate that FGF23 processing appears to be severely impaired in *Furin*^*osb*^*-/- mice* under conditions of iron deficiency as these mice maintained in the circulation a 1:1 ratio (i.e., 100%) of intact over C-terminal FGF23 and very low level of cleaved FGF23 (Figure 2G-H). Despite this near complete inhibition in FGF23 processing, iron deficient *Furin*^*osb*^*-/- mice* displayed a paradoxical increase in serum phosphate level compared to the control littermates fed the same diet (Figure 2I). Altogether these results suggest that furin maybe the main PC responsible of FGF23 processing under iron deficiency.

**Figure 2:**
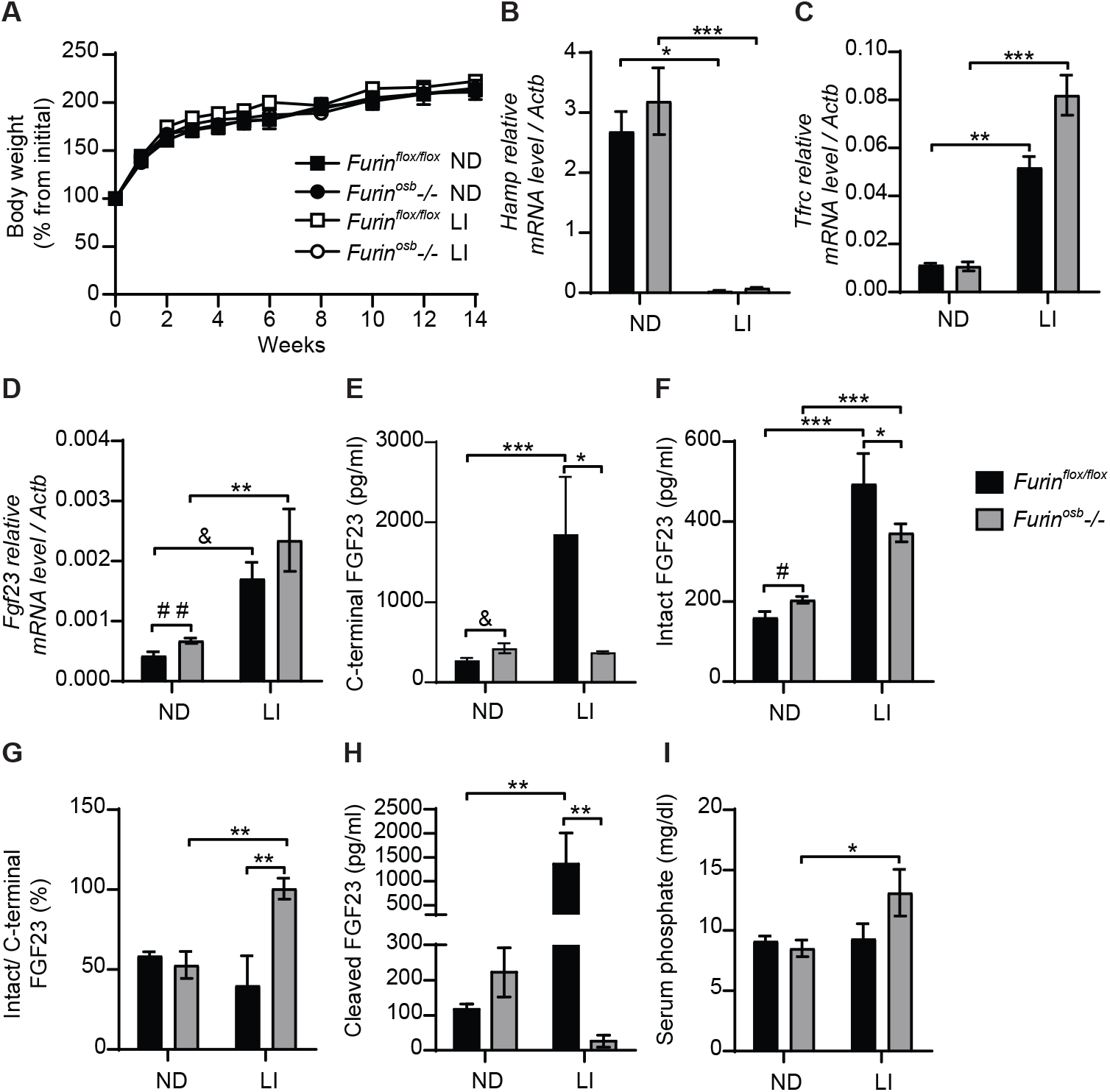
Impaired FGF23 processing in *Furin*^*osb*^*-/- mice* following iron restriction. *Furin*^*flox/flox*^ (n=3-5) and *Furin*^*osb*^*-/-* (n=7-8) 21-day-old mice were fed normal chow diet (ND) or a low iron diet (LI) for 14 weeks. **(A)** Body weight growth curve presented as percentage of initial body weight. **(B-C)** Expression of iron-regulated genes in liver. Hepcidin (*Hamp*) expression **(B)**. Transferrin receptor (*Tfrc*) expression **(C). (D)** *Fgf23* expression in long bone. **(E-H**) Plasma level of C-terminal FGF23 **(E)**, intact FGF23 **(F)**, percentage (%) of intact over C-terminal FGF23 **(G)** and cleaved FGF23 levels **(H)**, calculated as (C-terminal FGF23) - (intact FGF23). **(I)** Serum phosphate levels. Results represent the mean ± SEM. \**P* < 0.05, \*\**P* < 0.01, and \*\*\**P* < 0.001, by 2-way ANOVA with Bonferroni’s multiple comparisons test or by Student’s t-test (D and E) #: *P*<0.05; # #: *P*<0.01, # # #: *P*<0.001 &: 0.05<*P*<0.1.

### Furin does not process FGF23 in osteoblasts and osteocytes following rhEPO and IL-1β injections

Erythropoietin and interleukin 1 beta (IL-1β), which are increased in condition of anemia and inflammation respectively, are two potent inducers of *Fgf23* at the transcriptional level ^16–19,26,27^. In wild type mice, the majority of circulating FGF23 (i.e., >80%) is cleaved following injection of either recombinant human erythropoietin (rhEPO) or IL-1β ^16,17,19^, suggesting the action of one or several PCs. Hence, we tested if furin contributes to FGF23 processing after a single injection of rhEPO or IL-1β. We first confirmed that two relatively low doses of rhEPO (i.e., 300U/kg and 3000U/kg) were sufficient to induce in bones and bone marrow of wild type C57BL/6J mice the expression of two known EPO target genes, erythropoietin receptor (*Epor*) and erythroferrone (*Erfe*) (Figure 3A-D). These two doses also robustly induced *Fgf23* expression in both tissues (Figure 3E-F). We next assessed the effect of the same dose of rhEPO in control and *Furin*^*osb*^*-/- mice* on the various circulating forms of FGF23. Circulating levels of total (i.e., C-terminal) FGF23 were increased dose-dependently following rhEPO injections, while circulating intact FGF23 levels were only modestly changed in both control and *Furin*^*osb*^*-/- mice* (Figure 3G-H). The ratio of intact over C-terminal FGF23 level was robustly reduced by the same treatment (Figure 3I) and cleaved FGF23 was increased more than 10-fold by rhEPO in both control and *Furin*^*osb*^*-/- mice* (Figure 3J), suggesting that following rhEPO, FGF23 is normally processed even in absence of furin in osteoblasts and osteocytes.

**Figure 3:**
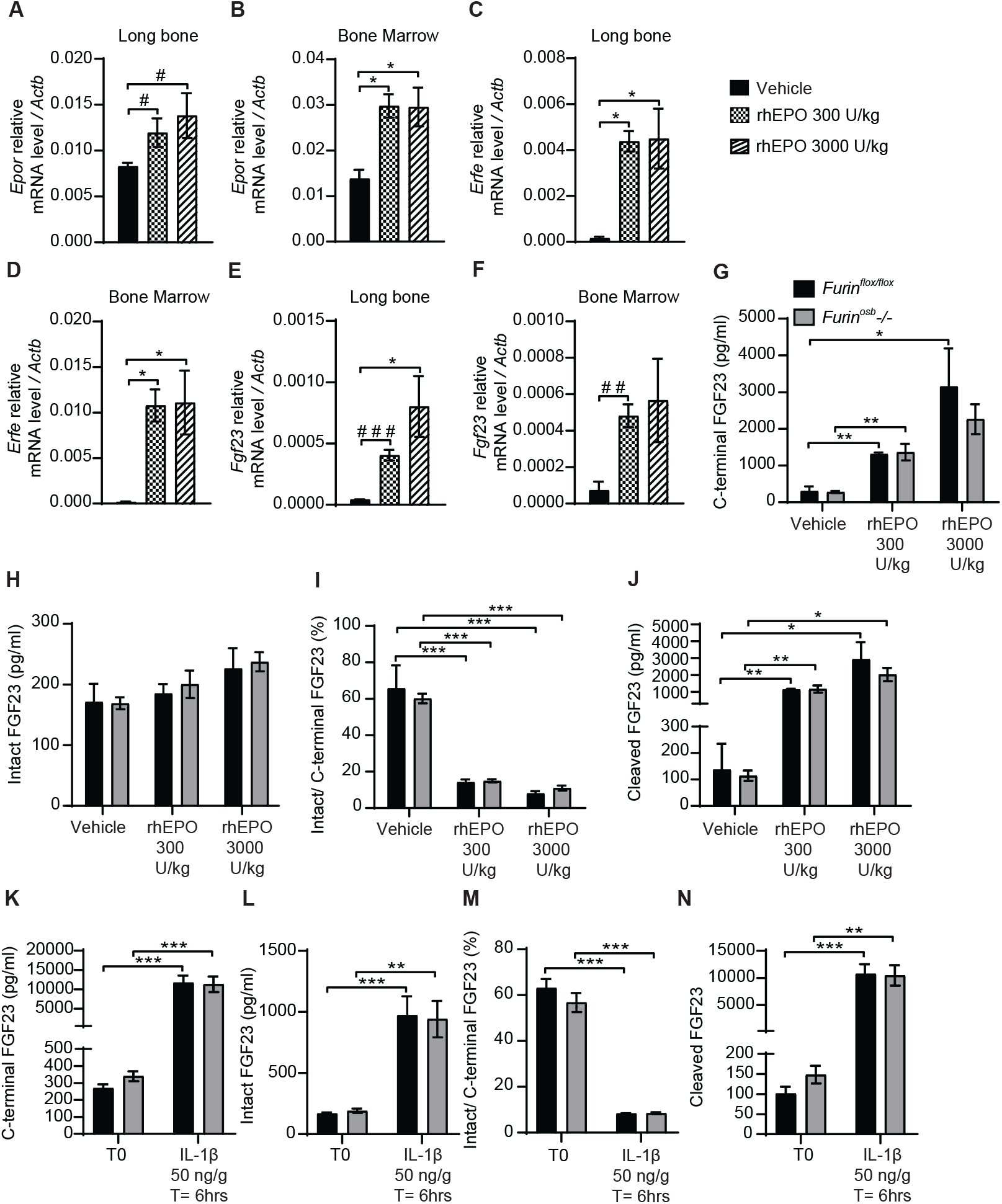
FGF23 is normally processed following erythropoietin and IL-1β injection in *Furin*^*osb*^*-/-*mice. Genes expression in long bone **(A, B and E)** and bone marrow **(C, D and F)** of 2-month-old C57BL/6J mice 6 hours following the injection of vehicle (n=4), 300 U/kg (n=4) or 3000 U/kg (n=4) of recombinant human erythropoietin (rhEPO). Erythropoietin receptor (*Epor*) **(A and C)**, erythroferrone (*Erfe*) **(B and D)** and FGF23 (*Fgf23*) **(C and F). (G-J)** Plasma FGF23 measurements in 4-month-old *Furin*^*flox/flox*^ (n=3) and *Furin*^*osb*^*-/-* (n=3) mice 6 hours following the injection of vehicle, 300 U/kg or 3000 U/kg of rhEPO. C-terminal FGF23 **(G)**, intact FGF23 **(H)**, percentage (%) of intact over C-terminal FGF23 **(I)** and cleaved FGF23 levels **(J)**, calculated as (C-terminal FGF23) - (intact FGF23). **(K-N)** Plasma FGF23 measurements in 4-month-old *Furin*^*flox/flox*^ (n=4) and *Furin*^*osb*^*-/-* (n=4) mice before (T0) and 6 hours following the injection of 50 ng/g IL-1β (T=6hrs). C-terminal FGF23 **(K)**, intact FGF23 **(L)**, percentage (%) of intact over C-terminal FGF23 **(M)** and cleaved FGF23 levels **(N)**. Results represent the mean ± SEM. \**P* < 0.05, \*\**P* < 0.01, and \*\*\**P* < 0.001, by 1-way ANOVA **(A-F)** or 2-way ANOVA **(G-N)** with Bonferroni’s multiple comparisons test, or by Student’s t-test **(A and E)** #: *P*<0.05; # #: *P*<0.01, # # #: *P*<0.001 &: 0.05<*P*<0.1.

In separate experiments, control and *Furin*^*osb*^*-/- mice* were administered a single dose of IL-1β (50ng/g), that induces *Fgf23* expression in bone and raises circulating total and cleaved FGF23 levels ^16^. Six hours after the injection of IL-1β, the circulating level of total (C-terminal) FGF23 increased over 30-fold, while intact FGF23 increased around 10-fold, in both control and *Furin*^*osb*^*-/- mice* (Figure 3K-L). Following the IL-1β treatment the ratio of intact over C-terminal FGF23 level decreased from 60% to less than 10% in both control and mutant mice (Figure 3M) and the plasma concentration of cleaved FGF23 increased more than 10 times (Figure 3N). Together, these data show that FGF23 processing occurs normally following rhEPO or IL-1β injections in the absence of furin in osteoblasts and osteocytes,

### Normal FGF23 processing in mice lacking furin in osteoblasts, osteocytes and hematopoietic cells following rhEPO injections

The data presented above suggest that FGF23 was normally processed following rhEPO or IL-1β injection in *Furin*^*osb*^*-/-*mice. Osteoblasts and osteocytes are the main source of circulating FGF23 in normal conditions ^27,28^, but bone marrow hematopoietic cells contribute to approximately 40% of circulating FGF23 following rhEPO treatment ^27^. Therefore, in *Furin*^*osb*^*-/-*mice, bone marrow hematopoietic cells could still be a significant source of cleaved FGF23 following rhEPO injections. To test this possibility, we generated mice lacking furin in all bone marrow hematopoietic cells (*Furin*^*BM*^*-/-*) by breeding *Furin*^*flox/flox*^ mice with *Vav1-Cre. Vav1-Cre* transgenic animals expresses the Cre recombinase under the control of the *Vav1* promoter, allowing the efficient deletion of furin in fetal and adult hematopoietic stem cells (HSC) ^29^ and hence in all hematopoietic cells. To address any potential redundancy between osteocytes and hematopoietic cells in the cleavage of FGF23, we also generated mice lacking *Furin* in both cell types, i.e., *Furin*^*flox/flox*^;hOCN-Cre;Vav1-Cre mice (*Furin*^*osb;BM*^*-/-*). Furin deficiency in T cells impairs the function of regulatory and effector T cells, causing at around 6 month of age severe inflammatory bowel disease, weight loss, ruffled hair and hunched appearance ^30^. Confirming efficient inactivation of *Furin* in hematopoietic cells, these defects were also observed in our *Furin*^*BM*^*-/- mice* between 4 and 6 months of age (data not shown). Hence, in the following experiments all mice were 8 weeks of age, an age when *Furin*^*BM*^*-/- mice* are still healthy.

A single injection of rhEPO (3000U/kg) increased circulating total (C-terminal) FGF23 by 20-30 fold and intact FGF23 by about 2 fold in *Furin*^*flox/flox*^, *Furin*^*osb*^*-/-, Furin*^*BM*^*-/-*and *Furin*^*osb;BM*^*-/-* mice (Figure 4A-B). The same treatment also reduced the ratio of intact over C-terminal FGF23 from 40% to <10%, while plasma cleaved FGF23 level increased to the same extent (>10 fold) in mice of all genotypes (Figure 4C-D). Together these results indicate that in vivo, FGF23 is still processed in absence of furin in osteoblasts/osteocytes and bone marrow hematopoietic cells in physiological condition and following rhEPO injections.

**Figure 4:**
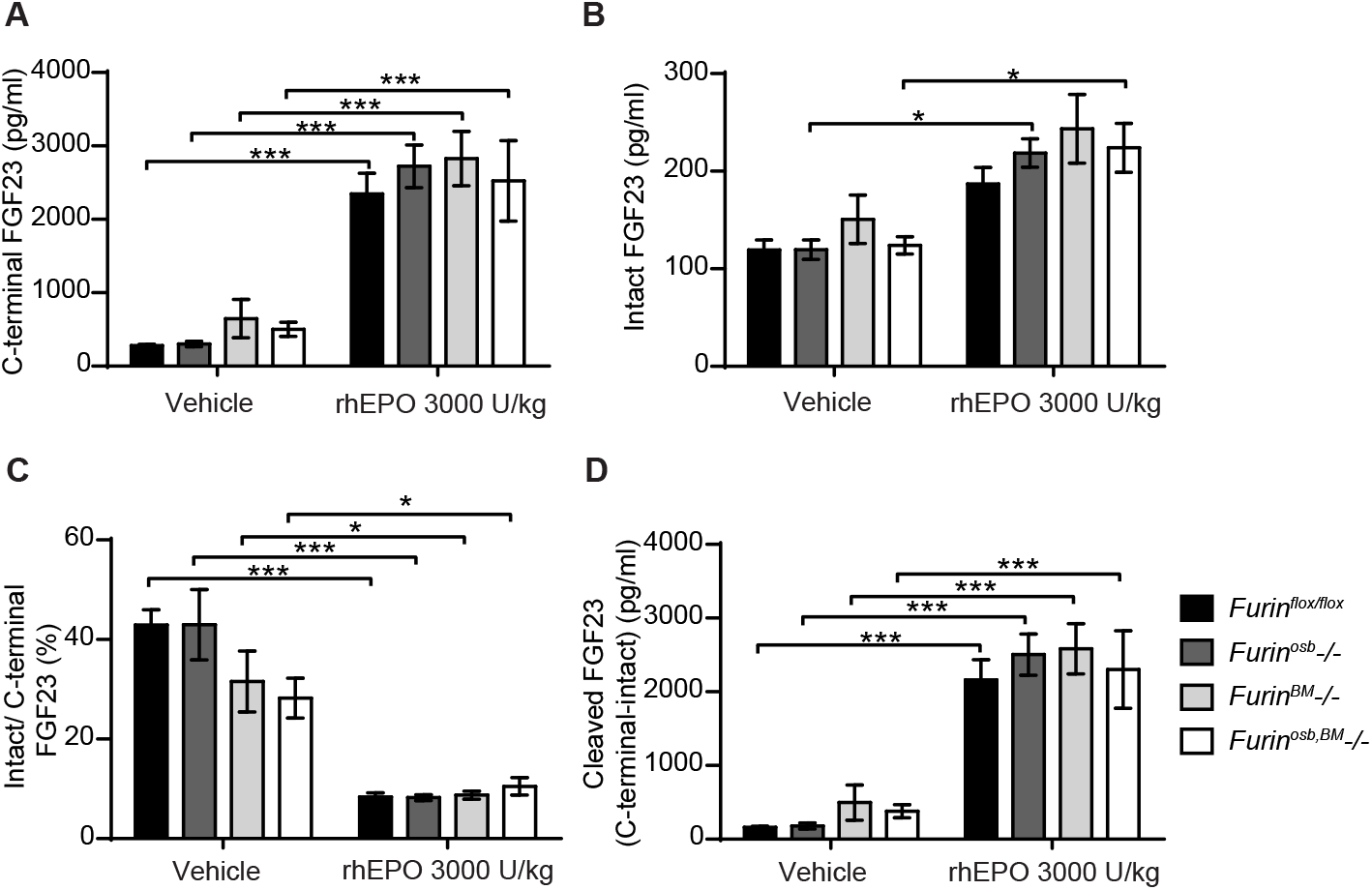
*Furin* inactivation in osteoblasts, osteocytes and hematopoietic cells did not impair FGF23 processing following erythropoietin injection. Plasma FGF23 measurements in 8-week-old *Furin*^*flox/flox*^ (n=7), *Furin*^*osb*^*-/-* (n=5), *Furin*^*osb*^*-/-* (n=5) and *Furin*^*osb;BM*^*-/-* (n=6) mice 6 hours following injection of vehicle or 3000 U/kg of recombinant human erythropoietin (rhEPO): C-terminal FGF23 **(A)**, intact FGF23 **(B)**, percentage (%) of intact over C-terminal FGF23 **(C)** and cleaved FGF23 levels **(D)**, calculated as (C-terminal FGF23) - (intact FGF23). Results represent the mean ± SEM. \**P* < 0.05 and \*\*\**P* < 0.001, by 2-way ANOVA with Bonferroni’s multiple comparisons test.

### Several PCs can process FGF23 in cell culture

Our data suggests that in some conditions (e.g., low phosphate diet, rhEPO injection or IL-1β injection), more than one PC may cleave FGF23 in vivo. In order to identify additional PC(s) that might process FGF23 in vivo, we tested FGF23 processing in cell lines expressing or not furin. In Chinese hamster ovary (CHO-K1) cells, which endogenously express furin ^31^, ectopically transfected human FGF23 is secreted and partially processed (Figure 5A). In the same cells, cleavage of FGF23 was inhibited by decanoyl-RVKR-CMK (RVKR) a pan-PC cell-permeable inhibitor, but not by hexa-D-arginine (D6R), a cell-surface PC inhibitor (Figure 5A). These observations suggest that cleavage of FGF23 occurs before it is secreted, in an intracellular secretory compartment, likely the trans-Golgi network ^32^ where furin, but also PC5A, PACE4 and PC7 are present. In addition, FGF23 was not cleaved in CHO-FD11 cells, which do not endogenously express furin ^31^, and co-transfection with a furin expressing vector was sufficient to restore efficient cleavage (Figure 5B, first two lanes). We next tested the capacity of other PCs, i.e., PC5A, PC5B, PACE4 and PC7 ^33^, to process FGF23 in absence of furin in the same cells. Among all these PCs, PC5A was the most efficient in processing FGF23 although PC5B, PACE4 and PC7 were also able to process FGF23 albeit less efficiently than PC5A (Figure 5B). Altogether, these results indicate that in addition to furin, several other PCs have the capacity to process FGF23 in vitro.

**Figure 5:**
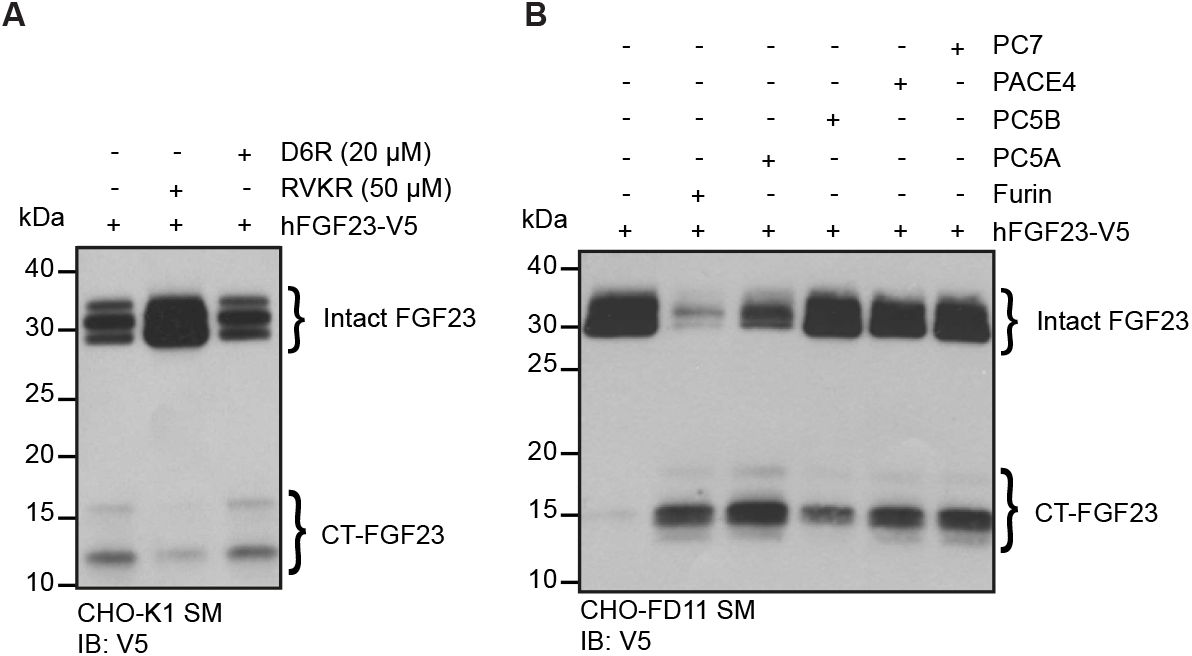
Multiple proprotein convertases can process FGF23 in mammalian cells. **(A)** Western blot analysis of the secretion media (SM) from CHO-K1 cells transfected with V5 tagged human FGF23 (hFGF23-V5) and cultured in the absence (-) or presence (+) of PC inhibitors, decanoyl-RVKR-CMK (RVKR) (50 μM) or hexa-D-arginine (D6R) (20 μM). **(B)** Western blot analysis of the secretion media (SM) from CHO furin-deficient FD11 cells co-transfected with V5 tagged human FGF23 (hFGF23-V5) and proprotein convertase Furin, PC5A, PC5B, PACE4 or PC7. Intact and processed C-terminal (CT) FGF23 forms are detected with V5 antibody.

### Genetic ablation of PC5 and furin in osteoblasts and osteocytes does not impair FGF23 processing

Since PC5A and PC5B, which are both encoded by *Pcsk5* ^33^, can cleave FGF23 in cell culture, and PC5A is expressed in differentiated osteoblasts and in osteocytes ^20,22^, we generated mice in which *Pcsk5* was inactivated specifically in osteoblasts and osteocytes by breeding *Pcsk5*^*fl/fl*^ mice ^34^ with hOCN-Cre transgenic animals. When fed a normal chow diet, control (*Pcsk5*^*fl/fl*^) and *Pcsk5*^*osb*^*-/-* mice displayed similar circulating level of intact FGF23 and serum phosphate (Figure 6A-B). Von Kossa/van Gieson staining on non-decalcified bone sections did not reveal any sign of osteomalacia in these mice (Figure 6C). These analyses suggest that PC5 in the osteoblast lineage is dispensable for the regulation of FGF23 and phosphate levels *in vivo*.

To test if PC5 and furin redundantly cleave FGF23 *in vivo*, we next generated mice lacking both furin and PC5 specifically in osteoblasts and osteocytes (i.e., *Furin*^*flox/flox*^; *Pcsk5*^*flox/flox*^; hOCN-Cre or *Furin;Pcsk5*^*osb*^*-/-*mice). Similar to the single inactivation of furin in the same cells, osteoblast and osteocytes-specific inactivation of furin and PC5 resulted in a modest increase in circulating intact FGF23 levels whether mice were fed normal or low phosphate diet (Figure 6D). Likewise, injection of rhEPO (300U/kg) increased circulating total (C-terminal) FGF23 by almost 10-fold and intact FGF23 by 25% in both control and *Furin;Pcsk5*^*osb*^*-/- mice* (Figure 6E-F). In these conditions, plasma intact over C-terminal ratio decreased by approximately 80%, while plasma cleaved FGF23 increased more than 10-fold, regardless of the mice genotype (Figure 6G-H). Altogether, these results show that FGF23 is still efficiently processed *in vivo* even when both furin and PC5 were inactivated in osteoblasts and osteocytes.

**Figure 6:**
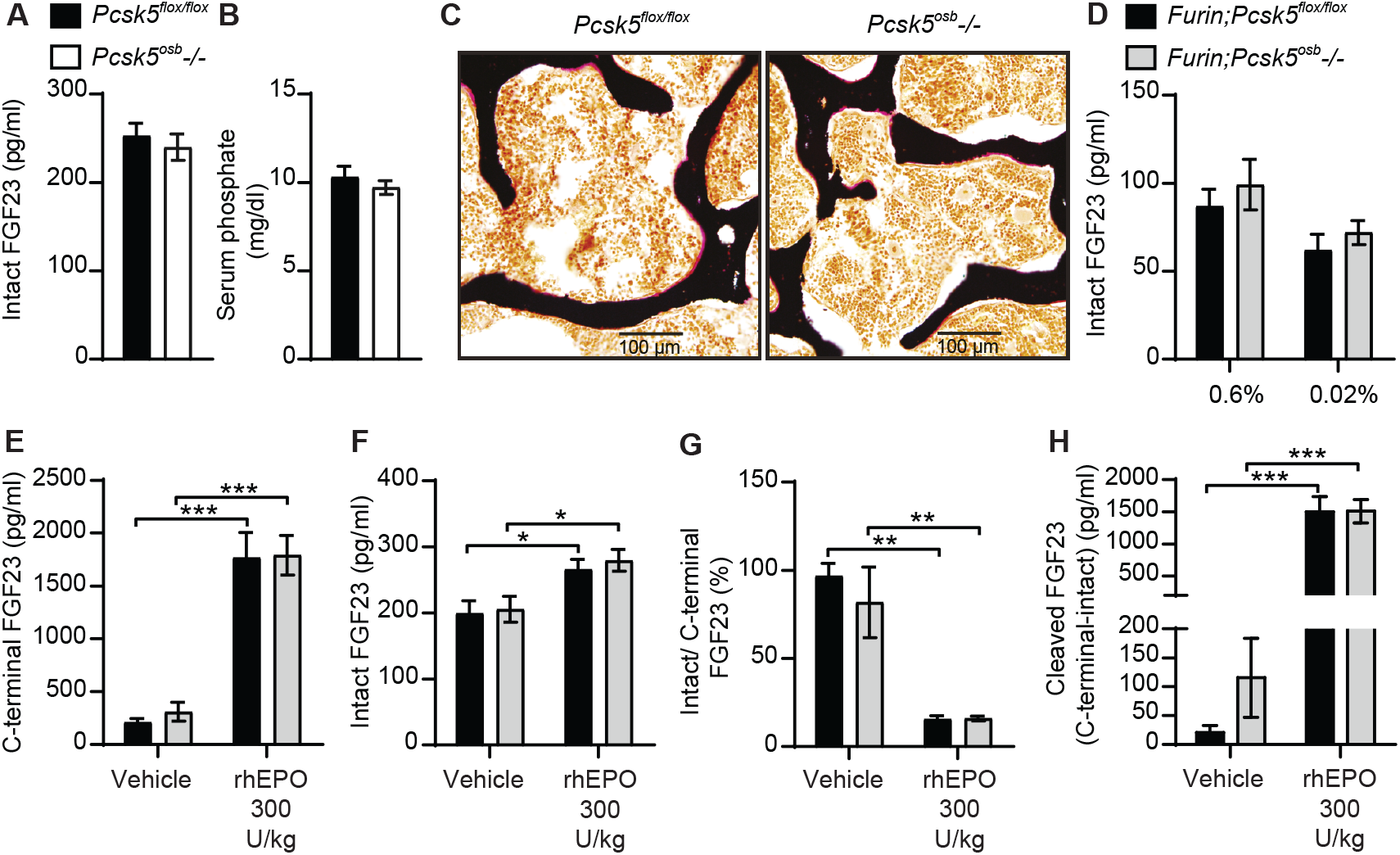
Genetic ablation of PC5 alone or in combination with Furin does not impair FGF23 processing *in vivo*. Plasma intact FGF23 **(A)** and serum phosphate levels **(B)** in 4-month-old *Pcsk5*^*flox/flox*^ (n=6) and *Pcsk5*^*osb*^*-/-* (n=7) mice fed a normal chow diet. **(C)** Von Kossa / van Gieson staining of vertebrae sections of *Pcsk5*^*flox/flox*^ and *Pcsk5*^*osb*^*-/-*. **(D)** Plasma intact FGF23 in 6-month-old *Furin;Pcsk5*^*flox/flox*^ (n=7-8) and *Furin;Pcsk5*^*osb*^*-/-* (n=8) fed a normal phosphate diet (0.6%) or a low phosphate diet (0.02%) for one week. **(E and H)** Plasma FGF23 measurements in 4-month-old *Furin;Pcsk5*^*flox/flox*^ (n=4) and *Furin;Pcsk5*^*osb*^*-/-* (n=4) 6 hours following an injection of vehicle and 300 U/kg of recombinant human erythropoietin (rhEPO): C-terminal FGF23 **(E)**, intact FGF23 **(F)**, percentage (%) of intact over C-terminal FGF23 **(G)** and cleaved FGF23 levels **(H)**, calculated as (C-terminal FGF23) - (intact FGF23). Results represent the mean ± SEM. \**P* < 0.05 and \*\*\**P* < 0.001, by 2-way ANOVA with Bonferroni’s multiple comparisons test, or by Student’s t-test **(A and B)**.

### Whole body PACE4 deletion does not impair FGF23 processing under physiological conditions or following IL-1β injections

In view of these negative results, we therefore asked whether PACE4 that is strongly expressed in mouse osteoblasts ^22^ and can cleave FGF23 *in vitro* (Figure 5B) does so in vivo. When fed a standard chow diet, *Pcsk6-/- mice* showed circulating levels of total (C-terminal) and intact FGF23 that were comparable to those of wild type littermates. Following a single IL-1β (50ng/g) injection, *Pcsk6-/- mice* displayed a significant decrease in circulating total (C-terminal) FGF23 while intact FGF23 was not significantly changed (Figure 7A and 7B). The ratio of intact over C-terminal FGF23 was equally decreased after IL-1β treatment in wild type and *Pcsk6-/- mice* plasma (Figure 7C). Circulating processed FGF23 was robustly increased by the same treatment regardless of the genotype, although PACE4 deficient mice did displayed a significant 30% reduction compared to wild type mice (Figure 7D). Altogether, these results show that PACE4 may partially contribute to FGF23 expression or secretion *in vivo* in response to IL-1β. Yet, FGF23 is still processed in absence of PACE4, suggesting the redundant action of other PCs.

**Figure 7:**
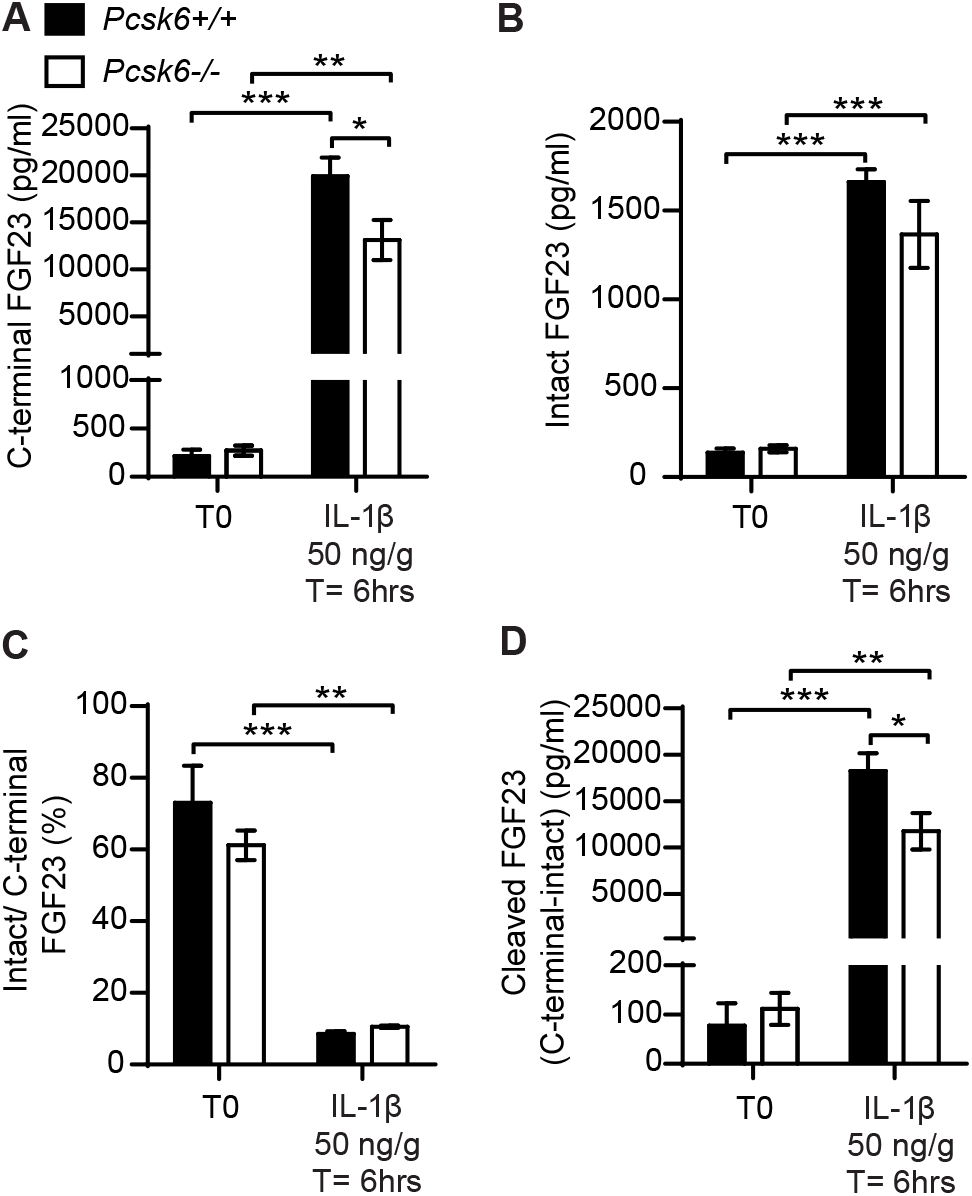
PACE4 inactivation in mice does not impair FGF23 processing. Plasma FGF23 measurements in 4-month-old *Pcsk6 +/+* (n=4) and *Pcsk6 -/-* (n=4) male mice before (T0) and 6 hours following the injection of 50 ng/g of IL-1β: C-terminal FGF23 **(A)**, intact FGF23 **(B)**, percentage (%) of intact over C-terminal FGF23 **(C)**, and cleaved FGF23 levels **(D)**, calculated as (C-terminal FGF23) - (intact FGF23). Results represent the mean ± SEM. \**P* < 0.05 and \*\*\**P* < 0.001, by 2-way ANOVA with Bonferroni’s multiple comparisons test.

## DISCUSSION

Over 20 years ago, mutations in a putative proprotein convertase (PC) target sequence within human FGF23 were found to cause ADHR in humans. Yet, despite the critical importance of this question for our understanding of phosphate metabolism, the identity of the specific PC(s) involved in FGF23 cleavage *in vivo* has remained elusive. We provide evidence here suggesting that furin, but not PC5, partially regulates FGF23 processing *in vivo* under normal conditions although this processing is by no means complete. The requirement of furin for FGF23 processing appears to be context or cell-type dependent, since FGF23 is still properly cleaved following rhEPO or IL-1β injection when furin is inactivated alone or in combination with PC5. Our results therefore suggest that additional yet to be characterized peptidases cleave FGF23 *in vivo*.

In mice fed standard chow diet, the modest increase in circulating intact FGF23 observed in *Furin*^*osb*^*-/- mice* did not cause hypophosphatemia and osteomalacia. This result is consistent with the observation that ADHR patients show variable age of onset and penetrance of the disease ^35^. Since iron deficiency positively correlates with intact FGF23 circulating levels in ADHR patient and results in hypophosphatemia and osteomalacia in mice carrying an AHDR mutation ^14,15^, we induced iron deficiency in control and furin deficient mice. Even under these conditions, furin deficient mice did not develop hypophosphatemia and in fact displayed higher serum phosphate level despite a near complete impairment of FGF23 cleavage. These observations suggest that although furin may be important for the cleavage of FGF23 in osteocytes in the context of iron deficiency, it may also be involved in the regulation of phosphate metabolism through FGF23-independent mechanism(s).

The results obtained in the context of iron deficiency raised the question as to whether, under normal conditions, FGF23 may also be cleaved by another PC that would be downregulated in osteocytes following iron deficiency. To clarify whether FGF23 cleavage by furin is context specific, we injected controls and furin-deficient mice with rhEPO or IL-1β. These two circulating factors are produced in response to anemia and inflammation respectively ^16,36^, and induce cleaved FGF23 secretion in bone and bone marrow cells in rodents ^16–19,26,27^. Herein, despite a strong increase in circulating total (C-terminal) FGF23 level following rhEPO or IL-1β injection, *Furin*^*osb*^*-/- mice* did not show a concurrent increase in intact FGF23 suggesting that either cleavage of FGF23 occurs by one or several other PC(s) in osteocytes, or that cleaved FGF23 originates mainly from bone marrow hematopoietic cells in this context. To address this second possibility, we generated mice deficient in furin in osteoblast-derived cells and/or in all hematopoietic cells. Our data shows that in these two cellular compartments, furin is dispensable for FGF23 cleavage following the injection of rhEPO further supporting the notion that furin is not the only peptidase involved in FGF23 processing *in vivo*.

PC5 (*Pcsk5*), PACE4 (*Pcsk6*) and PC7 (*Pcsk7*) are expressed in differentiated osteoblasts ^22^ and these PCs also target the minimal consensus cleavage site (i.e., RXXR) recognized by furin ^37^, suggesting that these enzymes may also cleave FGF23. Despite that, inactivation of *Pcsk5* specifically in osteoblasts and osteocytes did not impact intact FGF23 level and serum phosphate concentrations. In addition, mice deficient in both furin and PC5 in the osteoblast lineage are comparable to control mice in terms of FGF23 circulating level under physiological conditions and following rhEPO injection, suggesting that PC5 does not compensate for furin absence *in vivo*. Finally, even though PACE4 is highly expressed in differentiated osteoblasts ^22^ and cleaves FGF23 in mammalian cells, *Pcsk6* inactivation in mice only reduced FGF23 production following IL-1β injection and did not impact its cleavage.

Altogether, these findings provide genetic evidence that furin in osteoblasts and osteocytes is only partially responsible for FGF23 processing *in vivo*. Our data also showed that FGF23 cleavage is differentially regulated depending on the physiological context. Under conditions of iron deficiency furin is responsible for this process, while following single injection of rhEPO or IL-1β, additional proteases may be involved in FGF23 processing. Alternately, the increase in cleaved FGF23 following rhEPO and IL-1β injection could originate from non-osteoblastic cells such as bone marrow hematopoietic cells and involve multiple proteases including or not furin. Our study also excluded PC5 and PACE4 as major FGF23 processing enzymes *in vivo*. In conclusion, additional studies are required to determine precisely which enzyme(s) are involved in FGF23 cleavage *in vivo* and how furin regulates phosphate metabolism independently of FGF23.

## MATERIALS AND METHODS

### Mice and diets

*Furin*^*flox/flox*^ and *Pcsk5*^*flox/flox*^ mice were generated by introducing loxP site flanking exon 2 of furin gene and exon 1 of Pcsk5 gene as described previously ^38,39^. To generate conditional knockout of furin (*Furin*^*osb*^*-/-*) or PC5 (*Pcsk5*^*osb*^*-/-*) in osteoblasts and osteocytes, *Furin*^*flox/flox*^ and *Pcsk5*^*flox/flox*^ were crossed with OCN-Cre transgenic mice ^22^, which express the Cre-recombinase under the control of human osteocalcin promoter ^21^. *Furin;Pcsk5*^*osb*^*-/- mice* are deficient in both *Furin* and *Pcsk5* in osteoblasts/osteocytes and were generated by breeding *Furin*^*osb*^*-/-* and *Pcsk5*^*flox/flox*^ mice. *Furin*^*BM*^*-/-* and *Furin*^*osb;BM*^*-/- mice* were generated by breeding *Furin*^*osb*^*-/- mice* with *Furin*^*flox/+*^*;Vav-iCre* mice. The Vav-iCre expresses the Cre recombinase under the control of the *Vav* promoter, allowing the efficient deletion of furin in fetal and adult hematopoietic stem cells (HSC) ^29^. *Pcsk6-/- mice* were provided by Dr. Robert Day (Université de Sherbrooke), these mice were generated as described previously ^40^. All mice were backcrossed at least 10 time with the C57BL/6J genetic background, they were maintained at the IRCM pathogen free animal facility on 12h light/dark cycle and fed a normal chow diet. In all experiments, unless otherwise indicated, female mice were used.

In some indicated experiments mice were fed synthetic diet to induce iron or phosphate deficiency. To induce iron deficiency, after weaning at 21 days old, mice were fed either with normal chow diet or an iron deficient diet (Research diet, AIN-76A modified rodent diet, D09070102i) for 14 weeks in combination with bleeding equivalent to one blood capillary (70 μl) every two weeks. To induce phosphate deficiency, mice were fed for 5 days a normal phosphate diet (Envigo custom diet TD. 98243, 0.6% phosphorus), then divided to two groups. The first group was maintained for one more week on 0.6% PI diet while the second group was fed a phosphate deficient diet (Envigo custom diet, TD. 86128, 0.02% phosphorus). In other experiments, mice fed a normal chow diet were injected with vehicle (saline or PBS), 300U/kg or 3000U/kg of recombinant human erythropoietin (EPREX, Janssen-Cilag) or 50ng/g IL-1β (STEMCELL Technologies). All animal use complied with the guidelines of the Canadian Committee for Animal Protection and was approved by IRCM Animal Care Committee.

### Serum biochemistry

Intact and C-terminal FGF23 measurements were performed on EDTA plasma collected from mice under fed conditions using ELISA assay specific for mouse intact or C-terminal FGF23 (Quidel, cat#: 60-6800 and cat#: 60-6300). Since C-terminal FGF23 ELISA detects both intact and processed FGF23, it reflects the total concentration of FGF23 in the circulation. We also calculated the level of cleaved FGF23 by subtracting intact from C-terminal values. At end of experiments, mice were anesthetized, and serum was collected by heart puncture. Serum phosphate level was determined using a phosphorus assay kit (Sekisui Diagnostic, cat#:117-30). For urine phosphate measurements, urine was collected in the morning for 3 consecutive days and phosphate level was normalized to creatinine measured using creatinine assay (Quidel, cat#: 8009).

### RNA extraction

Liver, kidney and bone marrow RNA was extracted using the standard protocol previously described by Chomczynski *et al* ^41^, while bone RNA was extracted using Trizol reagent (Thermo Fisher). cDNA was generated by RNA reverse transcription using M-MLV Reverse Transcriptase (Thermo Fisher). Quantitative real-time PCR (qPCR) was performed using specific primers (Key Resources Table Appendix) and SYBR Green qPCR Master Mix (BiMake). Expression levels were normalized to *Actb* expression levels.

### Von Kossa/ Van Gieson staining

Mouse skeletons were collected and fixed in 10% formalin for 24 hours and in 70% ethanol for at least an additional 24 hours. Lumbar vertebrae were collected and dehydrated by gradual increase of ethanol percentage. Vertebrae were then embedded in methyl methacrylate resin as described previously ^42^. Vertebrae were sectioned at 7-µm sections and stained with Von kossa/Van Gieson and images were analyzed using the Osteomeasure analysis system (Osteometrics, Atlanta, GA).

### FGF23 processing in cell culture

The cDNAs coding for human furin, PC7, and PACE4, and mouse PC5A and PC5B were cloned into the pIRES2-EGFP vector ^43^, while the cDNA of hFGF23 was cloned in pIRES2-EGFP-V5. CHO-K1 and furin-deficient CHO-FD11 cells were cultured at 37 °C and 5% CO_2_ in Ham’s F-12 medium supplemented with 10% (v/v) FBS ^44^ and transfected with 3 μg of plasmid following the standard protocol for Lipofectamine 2000 (Thermo Fisher). For the PCs inhibition, CHO-K1 cells were transfected with cDNAs encoding for hFGF23-V5 and PCs in the ratio of 5:1 and 24 hours later pretreated for 5 hours with Dec-RVKR-CMK (50 μM; Tocris) or D6R (20 μM; Calbiochem) followed by 22 hours of treatment in serum free media. Cells supernatant was then collected in the presence of 1x complete protease inhibitor cocktail (Roche) and FGF23 processing was analyzed by SDS-PAGE (15% Tris-glycine) followed by western blot using V5 antibody (Thermo Fisher).

### Statistics

Statistical analyses were performed using GraphPad Prism software version 7.03. Results are shown as the mean ± SEM. Unpaired, 2-tailed Student’s *t* test was used for single measurements or comparison between two groups, while 1-way or 2-way ANOVA followed by Bonferroni’s post-test were used for comparison of more than 2 groups. A *P* value of less than 0.05 was considered statistically significant. All experiments were performed at least on 3 independent animals. Before starting each *in vivo* experiment, the mice were randomized into experimental groups with similar average body weight.

## ACKNOWLEDGEMENTS

We thank Dr. R. Day for providing *Pcsk6*^*-/-*^ mice, Dr. T. Clemens for sharing the hOC-Cre mice and Dr. T. Möröy for the iVav-Cre strain. This work was supported by funding from the Canada Research Chair program (M.F.) and the Natural Sciences and Engineering Research Council of Canada (RGPIN-2016-05213, to MF) and a CIHR Foundation grant (# 148363 to NGS). OAR received scholarships from IRCM and FRQS.

## Table Appendix

**Primers used**

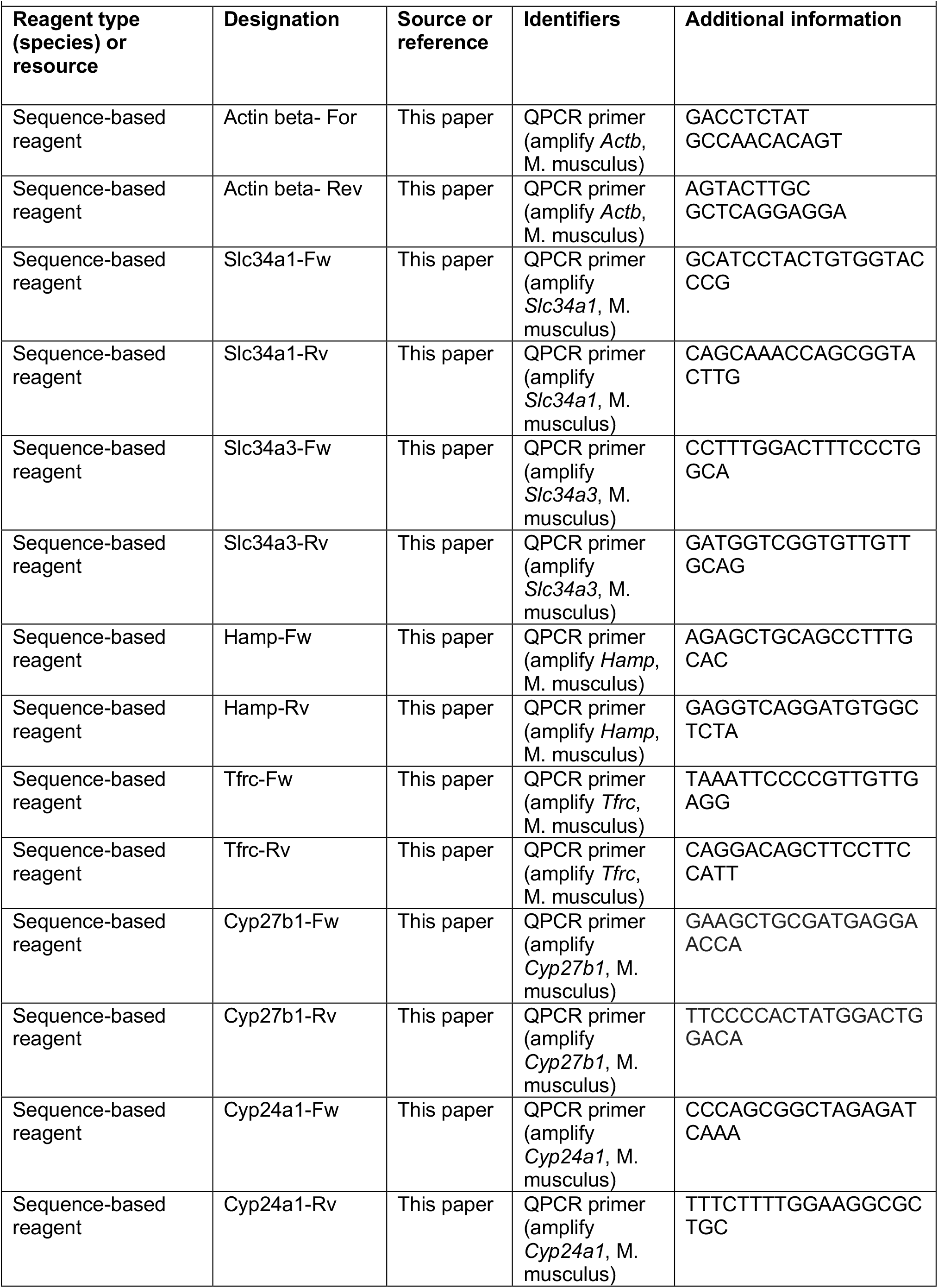
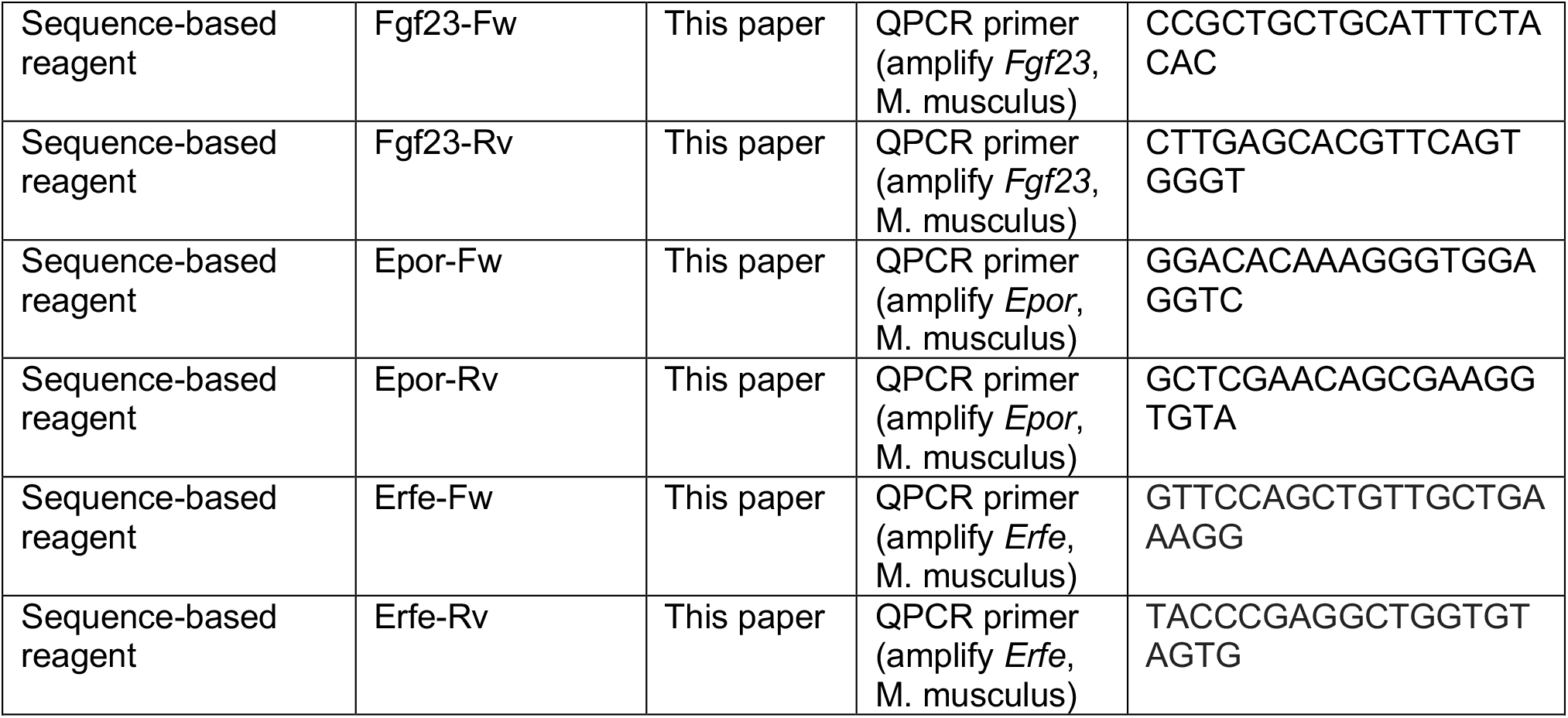

